# Nonsense mediated RNA decay factor UPF1 is critical for post-transcriptional and translational gene regulation in Arabidopsis

**DOI:** 10.1101/2020.03.02.971978

**Authors:** Vivek K. Raxwal, Craig G. Simpson, Jiradet Gloggnitzer, Juan Carlos Entinze, Wenbin Guo, Runxuan Zhang, John W.S. Brown, Karel Riha

## Abstract

Nonsense mediated RNA decay (NMD) is an evolutionary conserved RNA control mechanism that has also been implicated in the broader regulation of gene expression. Nevertheless, a role for NMD in genome regulation has not been fully assessed, partially because NMD inactivation is lethal in many organisms. Here, we performed in depth comparative analysis of Arabidopsis mutants lacking key proteins involved in different steps of NMD. We observed that UPF3, UPF1, and SMG7 have different impacts on NMD and the Arabidopsis transcriptome, with UPF1 having the biggest effect. Transcriptome assembly using stringent pipeline in UPF1-null plants revealed genome wide changes in alternative splicing, including switches in mRNA variants, suggesting a role for UPF1 in splicing. We further found that UPF1 inactivation leads to translational repression, manifested by a global shift in mRNAs from polysomes to monosomes and a downregulation of genes involved in translation and ribosome biogenesis. Despite this global change, NMD targets and low-expressed mRNAs with short half-lives were enriched in polysomes, indicating that UPF1 specifically suppresses the translation of aberrant RNAs. Particularly striking was an increase in the translation of TIR domain-containing, nucleotide-binding, leucine-rich repeat (TNL) immune receptors. The regulation of TNLs via UPF1/NMD-mediated mRNA stability and translational de-repression offers a dynamic mechanism for the rapid activation of TNLs in response to pathogen attack.

## Introduction

Accuracy and robustness of eukaryotic gene expression is controlled by several surveillance pathways. Nonsense mediated RNA decay (NMD) was discovered as a mechanism that inhibits translation of transcripts harboring a premature termination codon (PTC) (Peltz et al. 1993). Such aberrant RNAs can arise through alternative splicing, transcription errors, and missense mutations, and their translation would lead to dysfunctional, truncated proteins posing an unnecessary energetic burden to a cell. Studies across a range of organisms have revealed that NMD also affects a number of physiologically relevant transcripts and its activity is not constant, but is modulated in response to developmental and environmental cues (Karousis et al. 2016; Kim and Maquat 2019). This suggests a broader role for NMD in post-transcriptional gene regulation that goes beyond RNA surveillance.

We have gained fairly detailed insights into the molecular mechanism of NMD, particularly in yeast and humans (He and Jacobson 2015; Karousis et al. 2016; Kurosaki et al. 2019). In general, NMD consists of two steps: substrate recognition and substrate degradation. An essential event in substrate recognition is PTC definition. Depending on the organism, a PTC is defined contextually through the presence of an exon junction complex (EJC) deposited at splice junctions downstream of the termination codon and/or via a long 3’UTR. These RNA features promote accumulation of NMD factors that antagonize efficient termination of translation at a termination codon, leading to remodeling of the translating ribonucleoprotein particle and RNA decay (Kim and Maquat 2019). UPF1 is a highly conserved RNA helicase and ATPase that plays a central role in NMD. Although it binds promiscuously to any accessible part of RNA, it is particularly enriched at 3’UTRs (Hogg and Goff 2010; Hurt et al. 2013; Zund et al. 2013). In mammals, UPF1 and its kinase SMG1 form a complex with translation termination factors eRF1-eRF3 and out-compete termination-promoting factors, such as poly(A)-binding protein 1 (PABP). This can delay normal termination and initiate NMD. The key event in NMD substrate recognition is phosphorylation and activation of UPF1 by SMG1, which is triggered in the vicinity of an EJC by the associated factors UPF2 and UPF3 (Kashima et al. 2006). Another feature that promotes NMD is a long 3’UTR. It has been proposed that increasing the physical distance between a termination codon and PABP at the poly-A tail renders termination less efficient and enhances susceptibility to NMD (Amrani et al. 2004; Kertesz et al. 2006; Singh et al. 2008).

Once UPF1 is phosphorylated, it inhibits translation initiation and recruits factors mediating RNA decay (Isken et al. 2008). Two main degradation pathways that rely on phosphoserine binding proteins from the SMG5/6/7 family exist in humans. SMG6 is an RNA endonuclease that cleaves in the vicinity of a PTC and the cleaved ends are further degraded by the 5’-3’ exonuclease XRN1 and the exosome in 3’-5’ direction. SMG5 and SMG7 form a heterodimer that binds to phosphorylated UPF1 and recruits factors mediating deadenylation and decapping, which are followed by exonuclease RNA degradation (Unterholzner and Izaurralde 2004; Loh et al. 2013). The SMG6 and SMG5/7 pathways seem to be redundant as they act on similar RNAs (Colombo et al. 2017).

Although NMD is one of the best studied RNA processing pathways, we still do not fully comprehend its biological function(s) and impact on genome regulation. This is partially because inactivation of NMD is lethal in many model organisms, including Drosophila, mice, zebrafish, Arabidopsis, as well as in human cell lines (Medghalchi et al. 2001; Metzstein and Krasnow 2006; Yoine et al. 2006; Wittkopp et al. 2009; Hwang and Maquat 2011). This limits phenotypic analyses to cell cultures and hypomorphic mutants that retain residual NMD activity. Recently, Drosophila and human cells lacking essential NMD factors were rescued by inactivating a single apoptosis-promoting gene that was targeted by NMD (Nelson et al. 2016). This argues that deregulation of a specific gene or process, rather than a general failure in RNA surveillance, underly the lethality of NMD-deficient organisms.

In Arabidopsis, NMD dysfunction affects a numerous physiological processes, including circadian rhythms, auxin response, and ethylene signaling (Kwon et al. 2014; Merchante et al. 2015; Chiam et al. 2019), but its most pronounced impact is on pathogen defense. Mutants deficient in UPF1, UPF3, and SMG7 exhibit constitutive activation of the immune response, which, in the case of UPF1 inactivation, leads to lethality due to necrosis of germinating seedlings (Yoine et al. 2006; Jeong et al. 2011; Rayson et al. 2012; Riehs-Kearnan et al. 2012). The effect of NMD on the defense response appears to be physiologically relevant. In wild-type plants, NMD is downregulated in response to invading pathogenic bacteria, which leads to the activation of intracellular TIR domain-containing, nucleotide-binding, leucine-rich repeat (TNL) immune receptors through stabilization of their mRNAs (Gloggnitzer et al. 2014; Jung et al. 2020). The lethality of Arabidopsis UPF1-null mutants can be rescued by abrogating PAD4, which acts as pathogen signal transducer immediately downstream of TNLs (Riehs-Kearnan et al. 2012). Nevertheless, *upf1 pad4* mutants still exhibit severe developmental defects and growth retardation. This is in contrast to *upf3* and *smg7* mutants that display much milder growth phenotypes (Arciga-Reyes et al. 2006; Riehs et al. 2008). This observation can be rationalized in *upf3* mutants, which may still retain substantial NMD activity as plants utilize both EJC-dependent and long 3’UTR-dependent mechanisms for PTC definition (Kertesz et al. 2006; Kurihara et al. 2009; Kalyna et al. 2012; Drechsel et al. 2013; Degtiar et al. 2015). However, the phenotypic difference between *upf1* and *smg7* mutants is rather surprising. Plants do not contain orthologues of mammalian SMG5 and SMG6 and, therefore, degradation of NMD-targeted transcripts is expected to rely solely on SMG7. Thus, inactivation of SMG7 should fully abrogate NMD activity and result in phenotypes similar to UPF1.

In this study, we asked whether the phenotypic difference between Arabidopsis *upf3, smg7*, and *upf1* mutations can be attributed to their distinct impact on NMD. Furthermore, we took advantage of viable UPF1-null mutants to assess the full impact of this protein on post-transcriptional gene expression. Using genome-wide approaches we show that UPF1 has a profound effect on the transcriptome and splicing. UPF1 also promotes translation globally, while inhibiting translation of aberrant transcripts. In addition, we examined the role of UPF1 in regulating plant TNL immune receptors, whose activity must be strictly balanced to maintain levels sufficient for pathogen surveillance but below the full response required for pathogen defense. Our data indicate that UPF1 exerts its effect on plant immunity primarily through translational suppression of TNL receptors, which may provide a mechanism for tight regulation of threshold TNL activity.

## Results

### NMD factors have different impact on accumulation of PTC-containing (PTC+) transcripts

Arabidopsis mutants deficient in different NMD components exhibit distinct phenotypes. While UPF3 inactivation results in only mild growth abnormalities and no loss of fertility, UPF1-null plants die of massive necrosis shortly after germination (Arciga-Reyes et al. 2006; Yoine et al. 2006). SMG7-null plants are viable but show severe growth retardation and are infertile due to a defect in meiotic progression (Riehs et al. 2008; Capitao et al. 2018). This raised the question of whether the phenotypic variability observed in Arabidopsis NMD mutants reflects different degrees of NMD deficiency.

The lethality and severe growth retardation of *upf1-3* and *smg7-1* mutants, respectively, was rescued by inactivating PAD4 (Riehs-Kearnan et al. 2012). These rescued plants allowed us to compare the impact of SMG7 and UPF1 on NMD while minimizing secondary effects caused by autoimmunity. We examined NMD in mutants carrying T-DNA disruptions in conserved domains of UPF1 (*upf1-3 pad4*), SMG7 (*smg7-1 pad4*), and UPF3 (*upf3-1*) (Hori and Watanabe 2005; Arciga-Reyes et al. 2006; Riehs-Kearnan et al. 2012). We also included a weak *upf1-5* allele with an insertion in the 3’ UTR that decreases UPF1 expression (Arciga-Reyes et al. 2006). To assess NMD efficiency, we quantified the relative abundance of PTC+ and normal mRNAs by high resolution RT-PCR (Simpson et al. 2008). We selected 66 primer pairs from 59 genes that covered AS events (ASEs) that were previously shown to be affected by NMD deficiency (Kalyna et al. 2012) (**Supplemental Table 1**). Transcripts with at least a 5% change in abundance in mutant samples compared to control and a p-value ≤ 0.05 (Student’s T-test) were considered significant. Of the 254 transcripts we tested, 131 were significantly altered in at least one experimental condition (**Figure 1A**). Most of these transcripts (122) were changed in *upf1-3 pad4*, whereas UPF3 deficiency affected only 69 transcripts. Recognition of phosphorylated UPF1 by the SMG5/6/7 family of phosphoserine-binding proteins is one of the defining steps of NMD. In Arabidopsis, SMG7 is the only member of this protein family acting in NMD (Riehs et al. 2008; Benkovics et al. 2011; Capitao et al. 2018). Therefore, we expected loss of SMG7 to have a similar impact on NMD as UPF1. However, only 82 transcripts were affected in *smg7-1 pad4* compared to 122 in *upf1-3 pad4* (**Figure 1A**). Most transcripts altered in *upf3-1, upf1-5*, or *smg7 pad4* were also altered in *upf1 pad4* (**Figure 1B**) and the *upf1 pad4* plants exhibited the largest differences in individual transcript abundance compared to control (**Figure 1C**).

**Figure 1.**
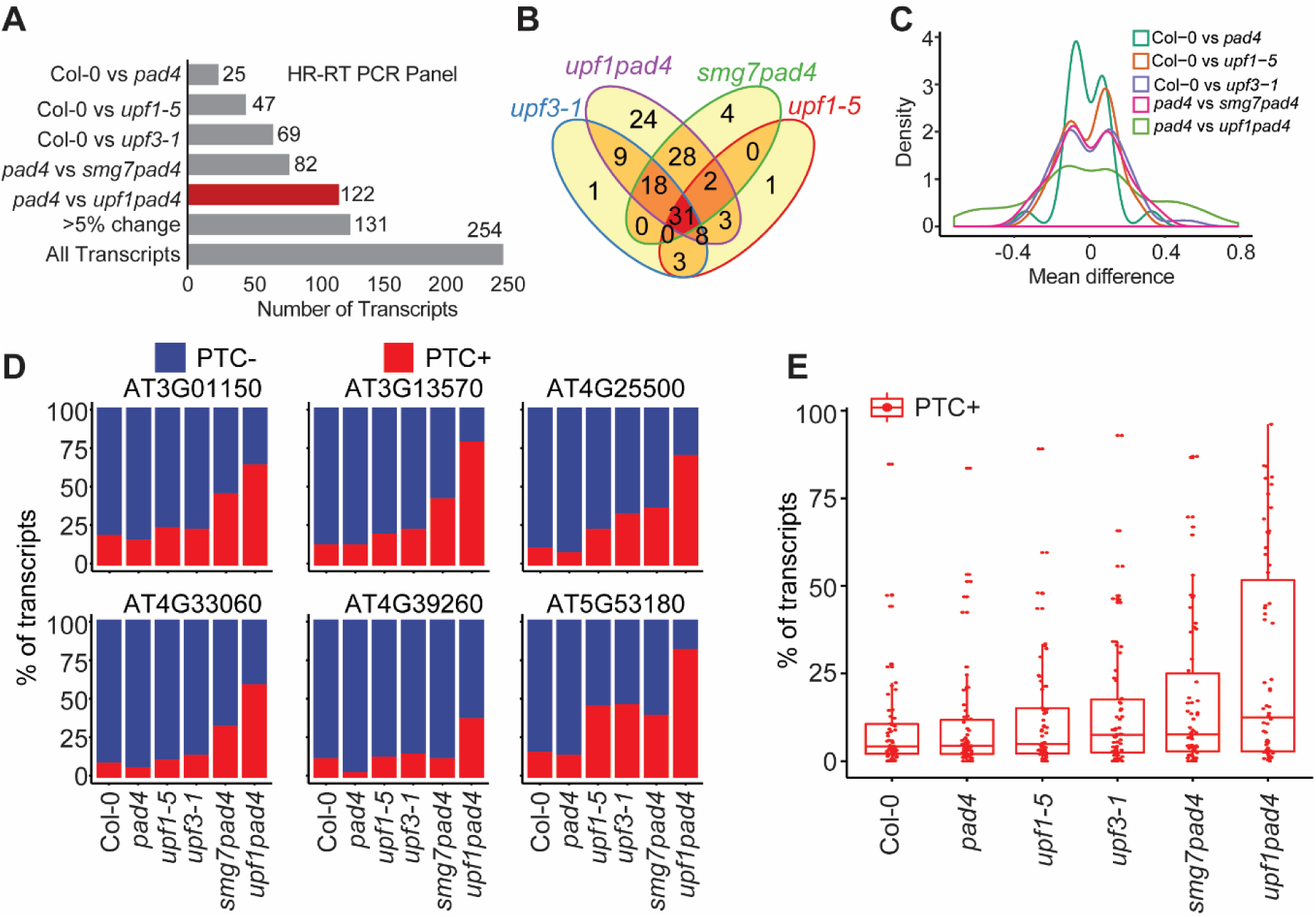
NMD efficiency in Arabidopsis NMD mutants scored using a high-resolution RT-PCR panel. (A) The distribution of transcript isoforms with significantly changed frequency in NMD mutants relative to control. (B) The overlap of transcripts with a significant change in abundance in NMD mutants. (C) A kernel density estimation plot showing the mean difference of transcript abundance in NMD mutant vs. wild type. (D) The percentage of PTC+ and normal transcripts (PTC-) for 6 ASEs in wild-type and different NMD mutants (AT3G01150 – PTB1; AT3G13570 – SCL30a; AT4G25500 – RS40; AT4G33060 – CYP57; AT4G39260 – RBGA6; AT5G53180 – PTB2). (E) Box plot showing the percentage distribution of PTC+ transcripts for the analyzed ASEs in different NMD mutants.

To assess whether the changes in transcript profiles can be attributed to NMD, we calculated the proportion of PTC+ transcripts for each ASE. As expected, PTC+ transcripts were relatively more abundant in NMD-deficient mutants than in wild type or *pad4* (**Figure 1D**,**E**). The different NMD mutations showed distinct impact on the frequency PTC+ transcripts for each ASE. For example, PTC transcripts from *AT3G13570* comprised 14% of transcripts in control samples (wild type or *pad4*), and their abundance increased to 24% in *upf3-1*, 44% in *smg7-1 pad4*, and 81% in *upf1-3 pad4* (**Figure 1D**). We observed a similar trend for *AT3G01150, AT4G25500*, and *AT4G33060*. In contrast, while *upf1-3 pad4* continued to show the strongest effect, we found a less pronounced effect of SMG7 than UPF3 in *AT5G53180* and *AT4G39260* (**Figure 1D**). From this analysis, we found that *upf1-3 pad4* had the strongest impact on PTC+ transcript accumulation, followed by *smg7-1 pad4, upf3-1*, and *upf1-5* (Fig. 1E). Indeed, *upf1-3 pad4* was significantly different from the other NMD mutants in this analysis (Mann-Whitney test, P < 0.05).

### UPF1 and SMG7 have distinct effects on transcriptome homeostasis

To further assess the effect of UPF1 and SMG7 on gene expression, we compared the leaf transcriptomes of *upf1 pad4* and *smg7 pad4* plants (Gloggnitzer et al. 2014) using RNA-seq. We found 17,393 and 15,933 genes to be expressed in the *pad4* vs. *upf1 pad4* and *pad4* vs. *smg7 pad4* contrast pairs, respectively [≥ 1 CPM; (counts per million) in at least two samples; **Supplemental Table 2**]. To compare the global impact of UPF1 and SMG7 deletions on the transcriptome, we plotted the kernel density distribution along with a scatter plot of gene expression between the two contrast pairs (**Figure 2A**). The wider distribution of the Gaussian curve in *upf1 pad4* samples (**Figure 2A**) indicates that UPF1 has a greater effect on transcriptional homeostasis than SMG7. Indeed, disruption of UPF1 affects the significant differential expression (DE) of 3,623 genes whereas only 612 were significantly altered in *smg7 pad4* mutants (log2 fold change ≥1 or ≤ −1, BH corrected p-value ≤0.01; **Fig 2B**). By comparing these datasets with the *upf3-1* transcriptome (Drechsel et al. 2013), we found that UPF3 has a lower effect on gene expression than SMG7 and UPF1 (**Supplemental Figure 1A**). NMD deficiency is expected to result in gene upregulation due to increased stability of aberrant RNAs. Indeed, more than 80% of the DE genes are upregulated in *smg7 pad4* and *upf3* mutants, but only 55% of DE genes are upregulated in *upf1 pad4* (**Figure 2B, Supplemental Figure 1B**), suggesting a broader deregulation of the *upf1 pad4* transcriptome.

**Figure 2.**
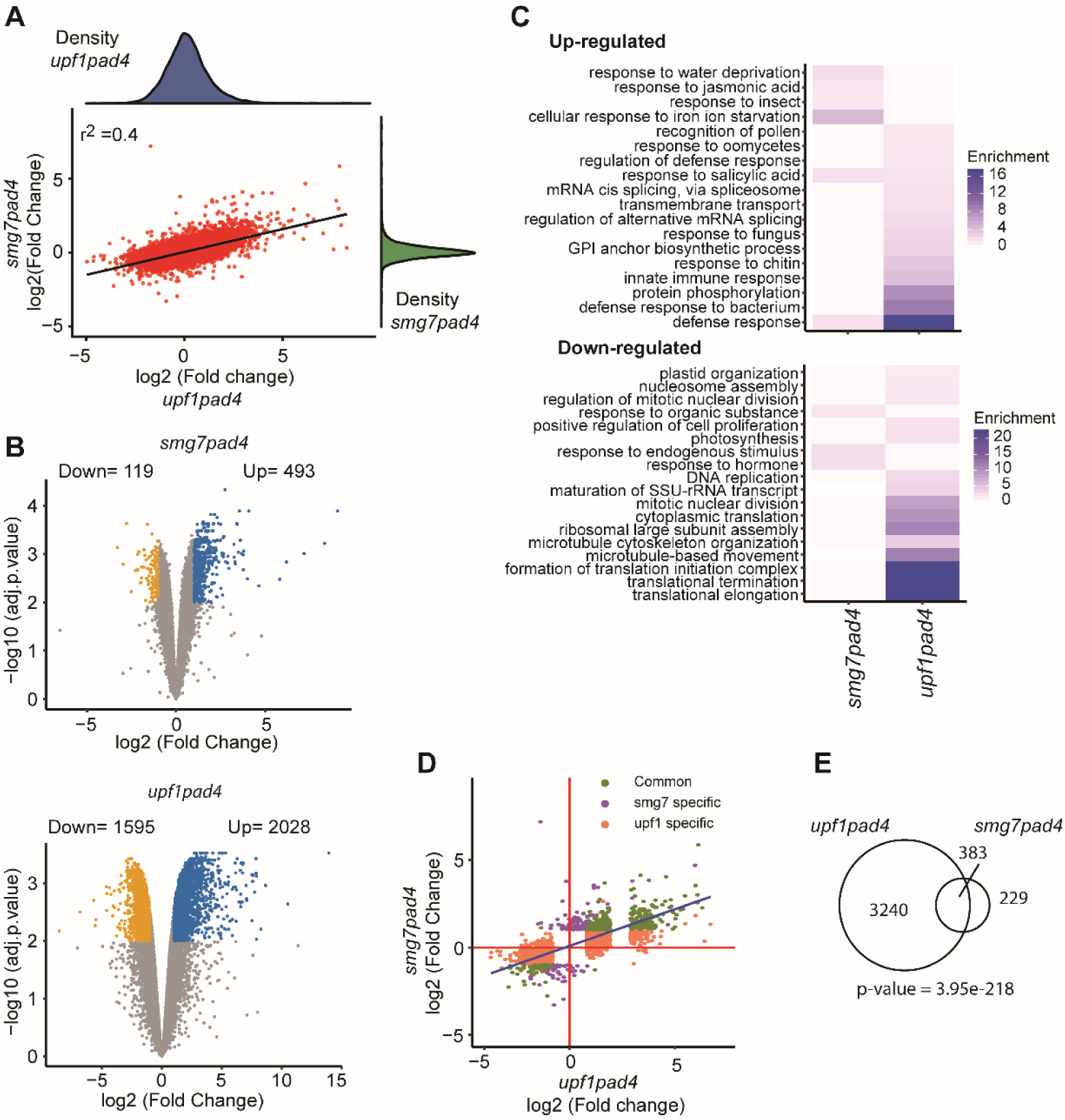
Transcriptome analysis of *upf1 pad4* and *smg7 pad4* mutants. (A) Scatterplot of gene expression profiles of *smg7 pad4* and *upf1 pad4*. Each dot represents log transformed mean expression of genes in *smg7 pad4* (Y-axis) and *upf1 pad4* (X-axis) compared to *pad4*. Smoothed kernel density estimates of gene expression are plotted along the X and Y axes (top and right, respectively). (B) Volcano plots depicting the distribution of significantly DE genes (BH corrected p-value ≤ 0-01 and log2(fold change) ≥ 1) in *upf1 pad4* and *smg7 pad4* as compared to *pad4*. (C) Enrichment (Fisher’s exact test, Bonferroni-corrected for P < 0.05) of gene ontological terms in *smg7 pad4* and *upf1 pad4* for up- and down-regulated genes. (D) Scatterplot of gene expression profiles in *smg7 pad4* and *upf1 pad4* with the distribution of DE genes specific to *upf1 pad4* or *smg7 pad4* and common to both. (E) An area proportional Venn diagram showing the overlap of DE genes between *upf1 pad4* and *smg7 pad4*. Statistical significance of the overlap was tested by Fischer’s exact test.

Gene ontological (GO) term enrichment analysis revealed that, despite abrogated pathogen signaling in *upf1 pad4* and *smg7 pad4* mutants, pathogen defense and responses to biotic stimuli are the main GO categories that are up-regulated in both mutants (**Figure 2C**). However, the number of GO categories related to plant defense and their degree of enrichment are substantially greater in *upf1 pad4* than in *smg7 pad4* plants. This may reflect the observation that, while PAD4 inactivation fully rescues growth in *smg7* mutants, *upf1 pad4* plants still exhibit retarded growth and abnormal development (Riehs-Kearnan et al. 2012). Another GO category up-regulated specifically in *upf1 pad4* was regulation of alternative mRNA splicing. UPF1, but not SMG7, also had a notable effect on processes related to translation and ribosome biogenesis, which were the most down-regulated GO categories in *upf1 pad4* (**Figure 2C**).

Next, we assessed the similarity between DE genes observed in *upf1 pad4* and *smg7 pad4*. More than 60% of *smg7 pad4* DE genes overlap with DE genes in *upf1 pad4* (hypergeometric test, p-value = 3.95e^-218^, **Figure 2D**,**E**). Together, our transcriptome data indicates that *smg7 pad4* DEs form a smaller subset of the *upf1pad4* DEs and that UPF1 has a much larger impact on the Arabidopsis transcriptome than SMG7 and UPF3.

### Arabidopsis NMD targets are processed through the decapping pathway

Whole-genome expression analysis in NMD mutants has been previously used to define NMD targets in Arabidopsis (Kurihara et al. 2009; Rayson et al. 2012; Drechsel et al. 2013). These studies used weaker NMD mutants which, together with their highly elevated pathogen response, partially impeded the identification of direct NMD targets. We took advantage of the *smg7 pad4* and *upf1 pad4* mutant plants, which have attenuated pathogen response and the strongest effect on the abundance of PTC-transcripts (**Figure 1**), to define NMD targets. A total of 333 DE genes were up-regulated in both *smg7 pad4* and *upf1 pad4* (**Figure 3A, Supplemental Table 2**), arguing that they constitute high-confidence NMD targets. Furthermore, the expression of these genes tended to be elevated in RNA-seq experiments of for *upf3, lba1* (a weaker allele of UPF1), *lba1 upf3*, and cycloheximide-treated plants (Drechsel et al. 2013), supporting the notion that this set of genes is enriched for canonical NMD targets (**Figure 3B**).

**Figure 3.**
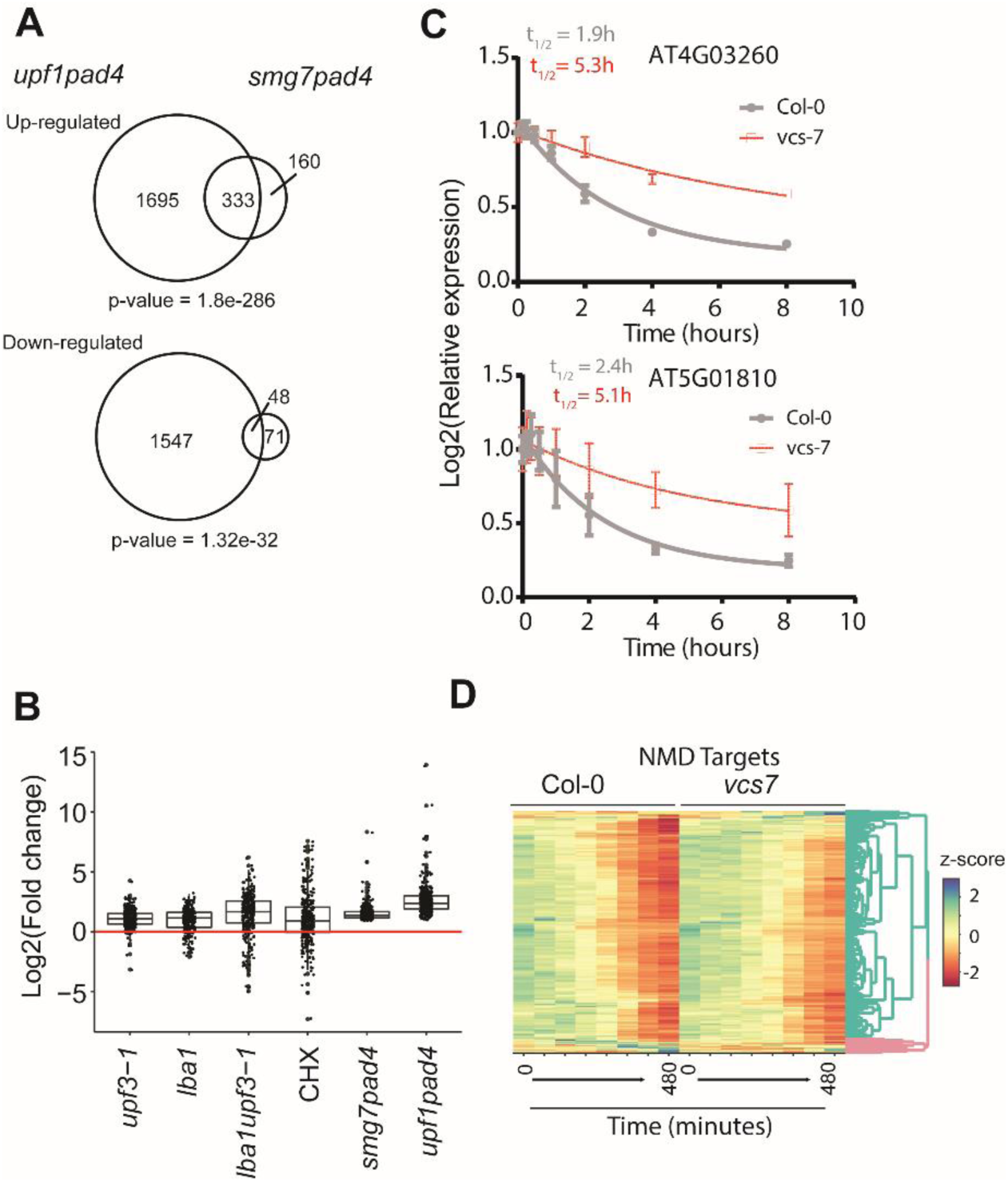
NMD targets are degraded by Varicose. (A) Area proportional Venn diagrams showing overlap between significantly DE genes of *smg7 pad4* and *upf1 pad4*. The statistical significance of the observed overlap was tested by Fischer’s exact test. (B) A box plot of a relative expression of NMD target genes in the indicated transcriptomes. (C) The decay rate of two representative genes in wild type and *vcs7*. (D) A clustered heatmap showing degradation kinetics of the 333 NMD target genes with time (0 to 480 minute) in wild type and *vcs7* after transcription inhibition.

NMD targets are expected to have decreased stability. Therefore, we explored recently published data on genome-wide stability of Arabidopsis RNAs in wild-type and *varicose* mutants (*vcs7*) to further characterize these 333 genes (Sorenson et al. 2018). Varicose is a scaffold protein that, together with DCP1 and DCP2, forms the decapping complex (Xu et al. 2006), a central component of the decapping-mediated RNA degradation pathway that contributes to RNA decay in Arabidopsis (Sorenson et al. 2018). We found that the vast majority of these genes (318 out of 333) show significantly reduced degradation in the *vcs7* mutant (**Figure 3C**,**D**) and, therefore, are likely bona fide NMD targets which undergo Varicose-dependent RNA decay.

### Transcript level analysis of UPF1-deficient plants from transcriptome reassembly

The high-resolution RT-PCR panel and RNA-seq analysis showed that UPF1 has the largest impact on aberrant RNAs and transcriptome homeostasis. To gain further insight into the effect of UPF1 on the Arabidopsis transcriptome, we made a detailed examination of gene expression in *upf1-3 pad4* at the level of individual transcripts. We firstly assembled transcripts from deep RNA-seq data obtained from wild type, *pad4*, and *upf1-3 pad4* seedlings and identified 1,601 transcripts that were not present in AtRTD2 (*Arabidopsis thaliana* reference transcript database) (Zhang et al. 2017). These transcripts were added to AtRTD2 and the new transcript reference was used to quantify transcript abundance and analyze differential gene and transcript expression. A total of 50,398 transcripts derived from 21,493 loci were expressed (> 1 CPM in at least one sample). 3,928 genes showed differential expression between *upf1 pad4* and *pad4* datasets that could be attributed to 4,542 transcripts (log2 fold change ≥1 or ≤ −1, BH corrected p-value <0.01, **Figure 4A**). Among these transcripts, 1,627 (36%) were down-regulated and 2,915 (64%) were up-regulated in *upf1 pad4* (**Supplemental Table 3**). 2,894 genes were DE only where the significant change in gene level expression is matched by the change in transcripts (**Figure 4A**).

**Figure 4.**
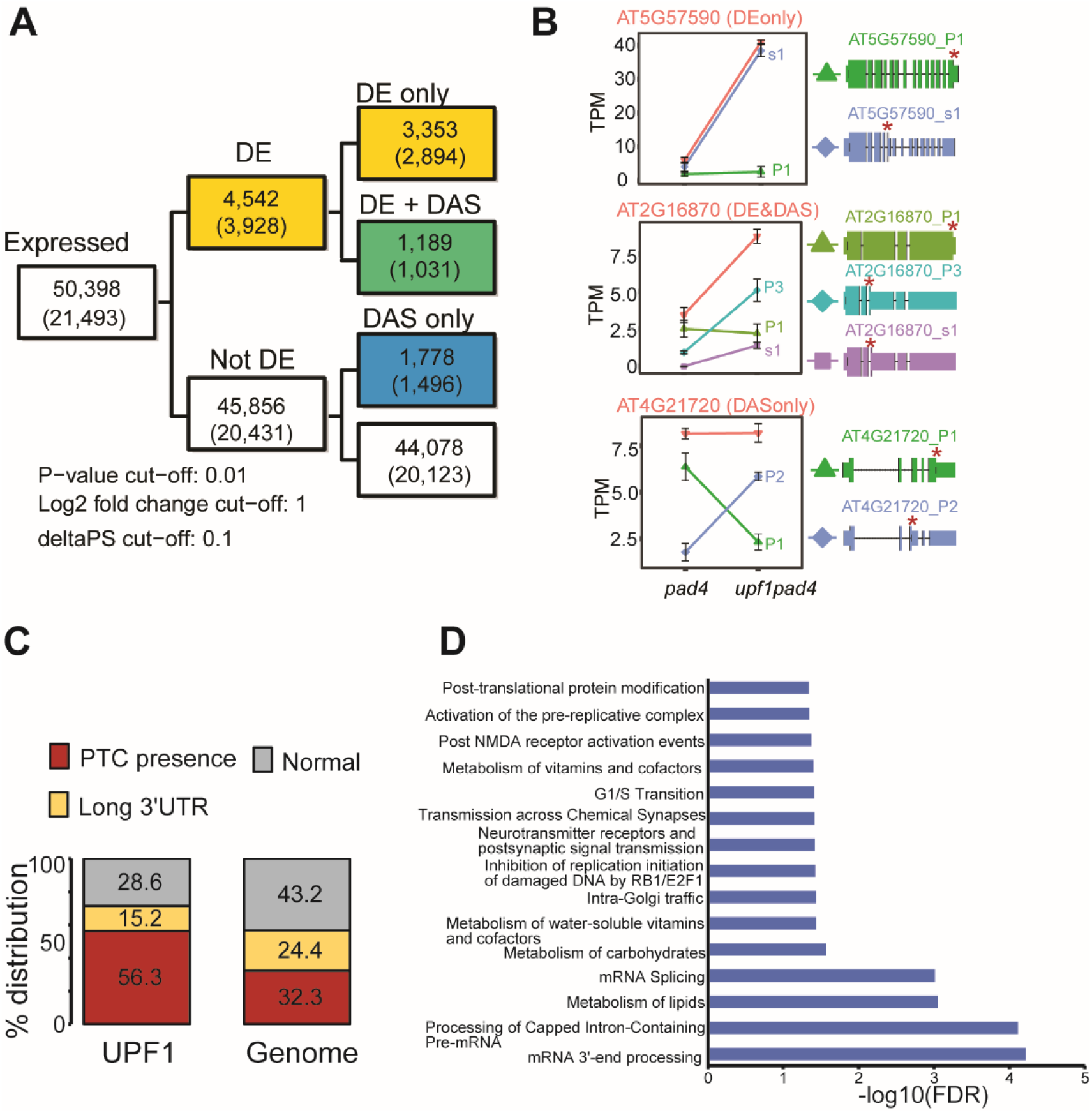
Transcript level analysis in *upf1 pad4* mutants. (A) Flow chart showing distribution of expressed transcripts and genes (numbers in brackets) in DE, DE+DAS and DAS gene categories. (B) Representative examples for DE (At5G57590), DE+DAS (At2G16870), and DAS (At4G21720) genes. Gene-level expression is represented in pink while other colors represent individual transcripts. Scaled representations of the transcript structure are shown with stop codons indicated (*). (C) The percentage distribution of PTC+ transcripts, normal transcripts, and transcripts with long 3’ UTRs for the combined transcripts identified in the DE only, DE and DAS, and DAS only categories. (D) Reactome pathway enrichment analysis (Fisher’s exact test, Bonferroni-corrected for P < 0.05) for combined transcripts in DE only, DE and DAS, and DAS only categories.

Some of these changes arose from the increased abundance of one or more transcripts from a locus, while other transcripts from the same loci remained unaltered (**Figure 4B**, *AT5G57590*). A significant portion of the transcripts upregulated in *upf1 pad4* are enriched for NMD features, such as PTCs defined by the presence of a splice site at least 50 bp downstream of a stop codon and/or a 3’UTR longer than 350 bp (Kalyna et al. 2012) (**Figure 4B**,**C**). This suggests that the increased transcript abundance is caused by stabilization of aberrant RNAs that are normally degraded by NMD.

Transcript-level analysis of DE genes also revealed a pattern consistent with differential alternative splicing (DE+DAS)(Calixto et al. 2018) at 1,031 loci. Typically in these cases, one alternatively-spliced isoform increased its abundance at the expense of other isoform (**Figure 4B**, *AT2G16870*). We also detected DAS only genes affecting 1,778 transcripts at 1,496 loci that did not exhibit overall significant changes in gene expression (**Figure 4A**,**B**, *AT4G21720*). This suggests an effect of UPF1 on splicing rather than just inefficient degradation of aberrant transcripts. In support of this notion, reactome pathway analysis of the loci with altered expression and splicing showed an enrichment for genes involved in mRNA processing and mRNA splicing (**Figure 4D**). Thus, UPF1 inactivation may affect splicing homeostasis by altering the expression of genes involved in splicing and RNA processing.

The transcriptome re-assembly identified 1601 novel RNAs that were not present in the AtRTD2 annotation of the Arabidopsis transcriptome (Zhang et al. 2017). These transcripts originated from either previously known genes (1,043) or novel loci (234). Examples for each of these cases are provided in **Supplemental Figure 2A**,**B.** The majority of the novel loci do not contain ORFs longer than 100 amino acids and were classified by Coding potential calculator (CPC2) as noncoding (**Supplemental Figure 2C**)(Kang et al. 2017). The expression of 8 transcripts from these novel loci was validated by qRT-PCR (**Supplemental Figure 2D**). Of these 1,601 novel transcripts, 1,341 were expressed at a level greater than 1 CPM in at least one sample and 459 were differentially expressed between *upf1 pad4* and *pad4* (BH corrected p-value ≤ 0.05, **Supplemental Table 4**). Among the 291 DE transcripts that were up-regulated in *upf1 pad4*, 70% contained a PTC and another 20% had a long 3’UTR. These transcripts originate from 228 previously annotated genes and 22 novel loci. The majority of these annotated genes were not significantly up-regulated in *smg7 pad4* (**Figure 5A**,**B**). Furthermore, this subset of transcripts tended to be stabilized in *vcs7* mutants (**Figure 5C**) indicating that they represent rapidly degraded RNAs that are processed by NMD in UPF1 proficient plants.

**Figure 5.**
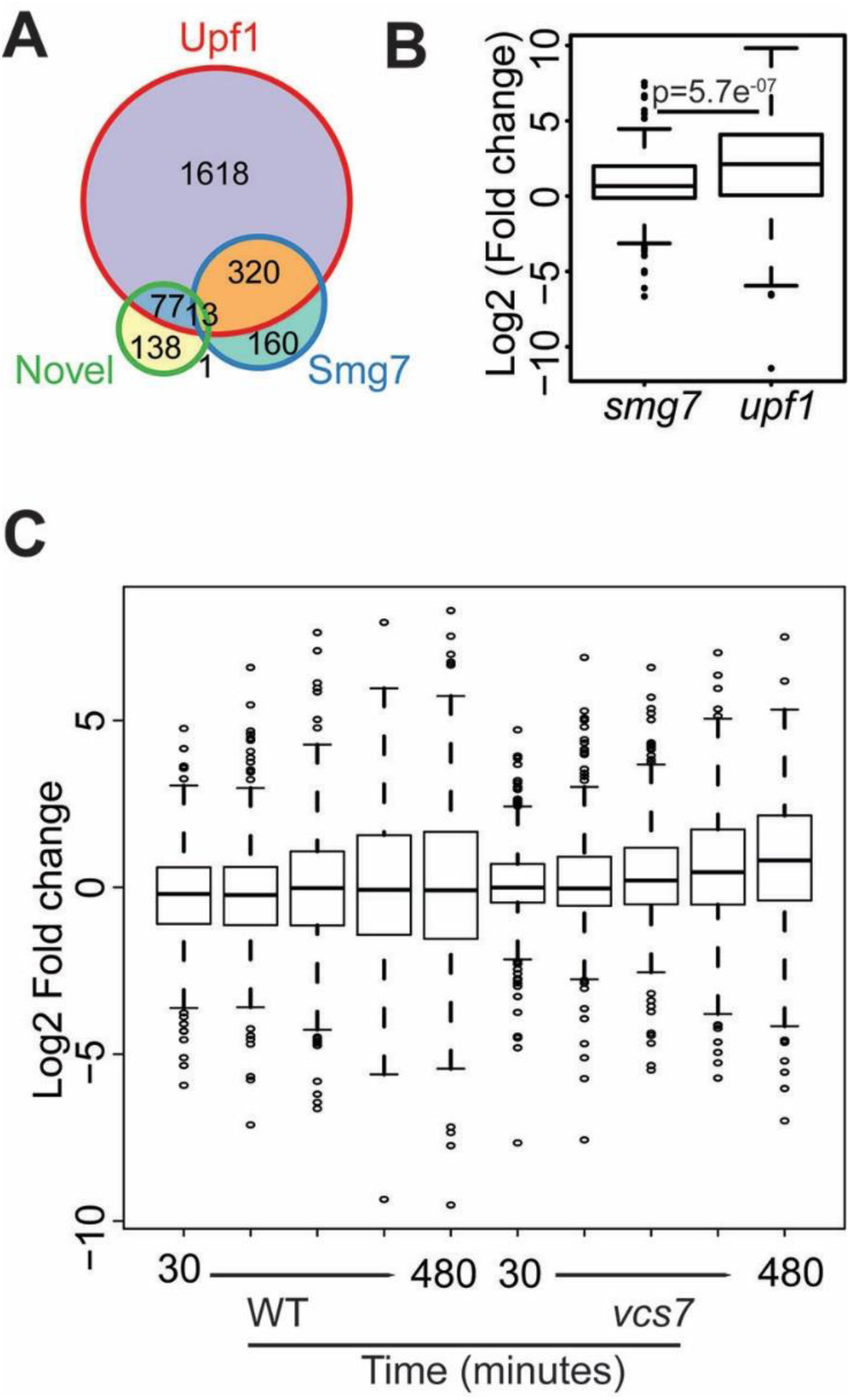
Expression of novel loci and gene isoforms upon UPF1 disruption. (A) Area proportional Venn diagram indicating the overlap of the 228 genes producing up-regulated novel transcripts from the assembly of *upf1 pad4* RNA-seq data (UPF1-specific) with genes upregulated in *smg7 pad4* and *upf1 pad4*. (B) Box plot showing the relative expression (Log2 fold change to *pad4*) of UPF1 specific transcripts in *smg7 pad4* and *upf1 pad4*. Statistical significance for differences in transcript expression was tested using Wilcoxon Signed-rank test. (C) Boxplot showing the degradation kinetics of UPF1 specific transcripts with time (30 to 480 minute) in wild type and *vcs7* after inhibiting transcription.

### The impact of UPF1 on translation

The pronounced down-regulation of GO categories related to translation and ribosome biogenesis in *upf1 pad4* plants (**Figure 2C**) prompted us to examine the global impact of UPF1 on the translatome. One of the established approaches for assessing translation is calculating the translation index, a ratio of the polysomal to cytoplasmic fraction of a given RNA. However, this method may produce skewed data as not all RNA molecules detected in the cellular transcriptome associate with cytoplasmic ribosomes (Larsson et al. 2010). A recent study in yeast demonstrated that RNAs contained in 80S monosomes undergo active translation and that most mRNAs exhibit some degree of monosome occupancy (Heyer and Moore 2016). Furthermore, since the detection of NMD-substrates inhibits further translation, NMD targets are predominately found on monosomes (Isken et al. 2008; Heyer and Moore 2016). Therefore, we decided to analyze the impact of UPF1 deficiency on the distribution of RNAs in monosomes and polysomes, rather than calculating the translation index.

Sequencing RNA from monosomal and polysomal fractions of *pad4* and *upf1 pad4* showed a strong linear correlation in the distribution of transcripts between monosomes and polysomes (**Figure 6A**, *pad4* r^2^ = 0.92, *upf1pad4* r^2^ = 0.88, p-value <2.2e-16). This indicates that the majority of mRNAs are proportionally distributed in both fractions. To look in more detail at the global effect of UPF1 loss on mRNA distribution between monosomes and polysomes, we calculated the relative polysome abundance for each gene in *upf1 pad4* and in *pad4* mutants [**Figure 6B**; expression in polysome (TPM)/expression in polysome + monosome (TPM)]. While the majority of transcripts are slightly more abundant on polysomes than monosomes in *pad4*, we found that there is a significant, global shift to the monosomal fraction in *upf1 pad4* (p-value <2.2e-16, KS test) (**Figure 6B**).

**Figure 6.**
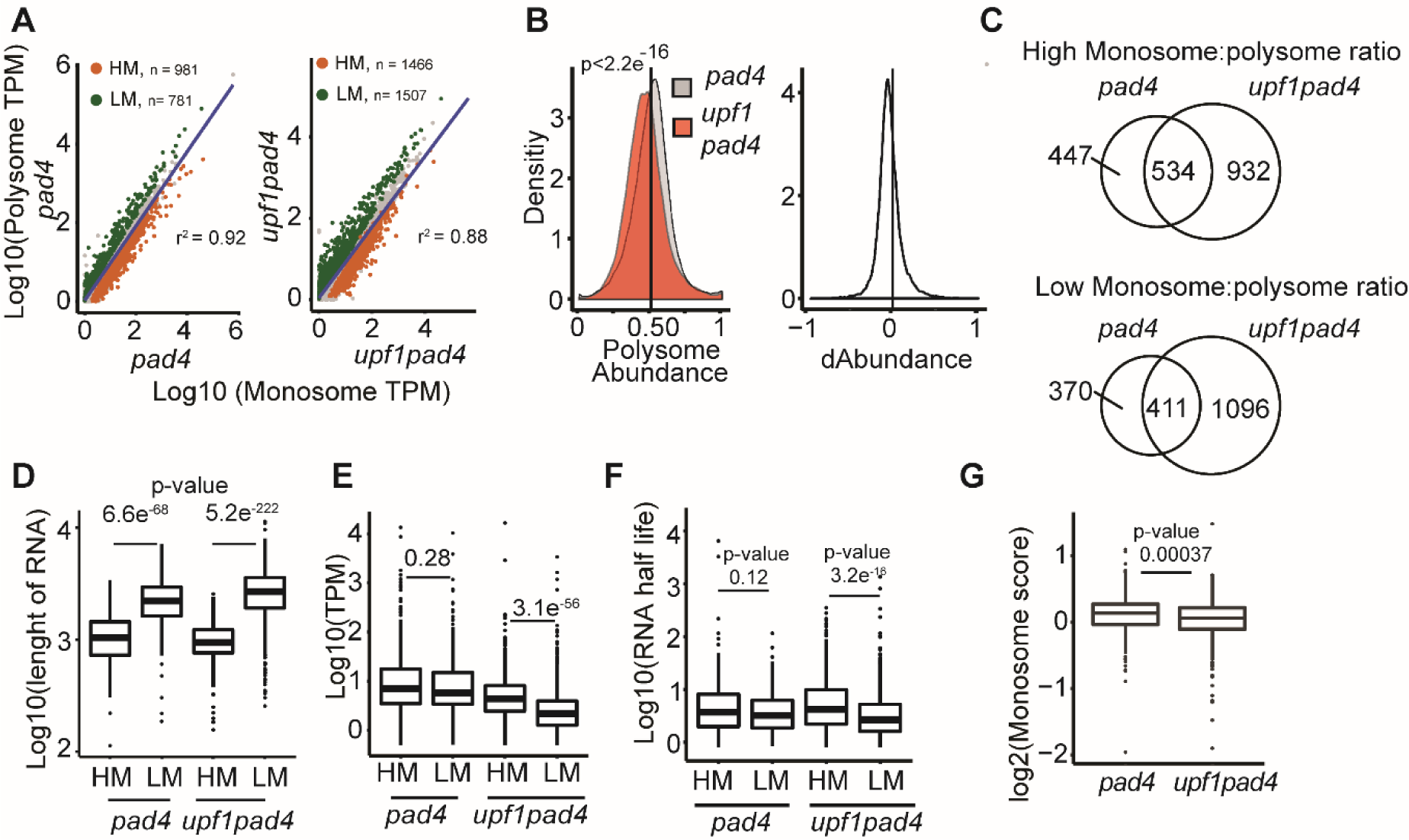
Translatome analysis in *upf1 pad4* mutants. (A) A scatter plot comparing gene abundance in monosome (X-axis) and polysome (Y-axis) fractions of *pad4* and *upf1 pad4*. The numbers of genes exhibiting high monosome (HM) or low monosome (LM) occupancy are indicated. B) A kernel density plot showing the distribution of genes according to their polysome abundance (x-axis) in *pad4* and *upf1 pad4*. Statistical significance for difference in the density plots was tested using Kolmogorov-Smirnov test. (C) Area proportional Venn diagram depicting overlap between HM and LM genes in *pad4* and *upf1 pad4*. The statistical significance of the observed overlap was tested by Fischer’s exact test. (D-F) Box plots showing (D) difference in gene length, (E) steady state transcription, and (F) half-life of HM and LM in *pad4* and *upf1 pad4*. Statistical significance for differences between HM and LM genes was tested using the Wilcoxon Signed-rank test. (G) Box plot depicting fold change in monosome score for NMD targets. Statistical significance was tested using Wilcoxon Signed-rank test.

To analyze genes that deviate from the proportional distribution between monosomes and polysomes, we calculated the monosome:polysome score as a fold enrichment between monosomal and polysomal bound RNA [Expression in monosome (TPM)/ Expression of in polysome (TPM)] and determined the differential distribution of RNAs in the monosomal and polysomal fractions using the limma-voom pipeline (Law et al. 2014). A total of 1,732 genes in *pad4* significantly deviated in their monosome:polysome score (BH corrected p-value < 0.05) with 981 genes (56 %) having a higher monosome occupancy (**Figure 6A**). In the absence of UPF1, 2,881 genes showed a significant deviation, with 1,466 genes (49%) displaying a higher monosome occupancy (**Figure 6B**). A substantial overlap (55%) in genes with either a high or low monosome:polysome score in both *pad4* and *upf1pad4* samples indicates a general tendency for these genes to associate with either monosomes or polysomes (**Figure 6C, Supplemental Table 5**).

We next compared the features of genes that exhibit high or low monosome occupancy in either *upf1 pad4* or *pad4* plants. As expected, we found that high monosome/low polysome occupancy is typical for shorter transcripts in both *upf1 pad4* and *pad4* (**Figure 6D**). While there is no difference in average gene expression levels between genes with low and high monosome occupancy in *pad4* plants, genes with low monosome/high polysome occupancy tend to show lower expression in *upf1 pad4* mutants (**Figure 6E**). We also noticed that this fraction is enriched for less stable transcripts (**Figure 6F**). Thus, inactivation of UPF1 leads to increased polysomal occupancy, and hence to increased translation of rarer and less stable short RNAs. To determine whether UPF1 inhibits translation of putative NMD targets, we quantified the monosome:polysome score in the 333 putative NMD targets (**Figure 3A, Supplemental Table 2**). Indeed, we observed increased polysome occupancy in these genes in *upf1 pad4* compared to *pad4* (**Figure 6G**). Together, these data indicate that while UPF1 inactivation leads to a general shift of transcripts to monosomes, presumably reflecting the global decrease in translation, it also increases polysomal abundance, and hence translation, of rare, less stable transcripts and putative NMD targets.

### UPF1 represses translation of TNL immune receptors

Previously, we showed that SMG7 regulates plant defense responses by controlling the turnover of TNL mRNAs (Gloggnitzer et al. 2014). The strong autoimmune response of *upf1* prompted us to ask whether TNLs are also targeted by UPF1. Comparing the transcriptomes of *upf1 pad4* and *smg7 pad4* we observed a significant (BH-corrected p-value ≤ 0.01and log2foldchange ≥ 1) up-regulation of 37 and 22 TNLs in *upf1 pad4* and *smg7 pad4*, respectively (**Figure 7A**). While 18 TNLs were upregulated in both datasets, 19 were exclusively increased in *upf1 pad4*. This is in agreement with UPF1’s stronger impact on the transcriptome as well as the strong deregulation of defense response genes in *upf1 pad4* plants (**Figure 2C**). Based on their stabilization in the absence of Varicose (**Figure 7B**), similar to NMD transcripts, the majority of NMD-regulated TNLs are subject to decapping-mediated 5’-3’ exonucleolytic degradation. This reinforces the view that TNLs are targets of NMD.

**Figure 7.**
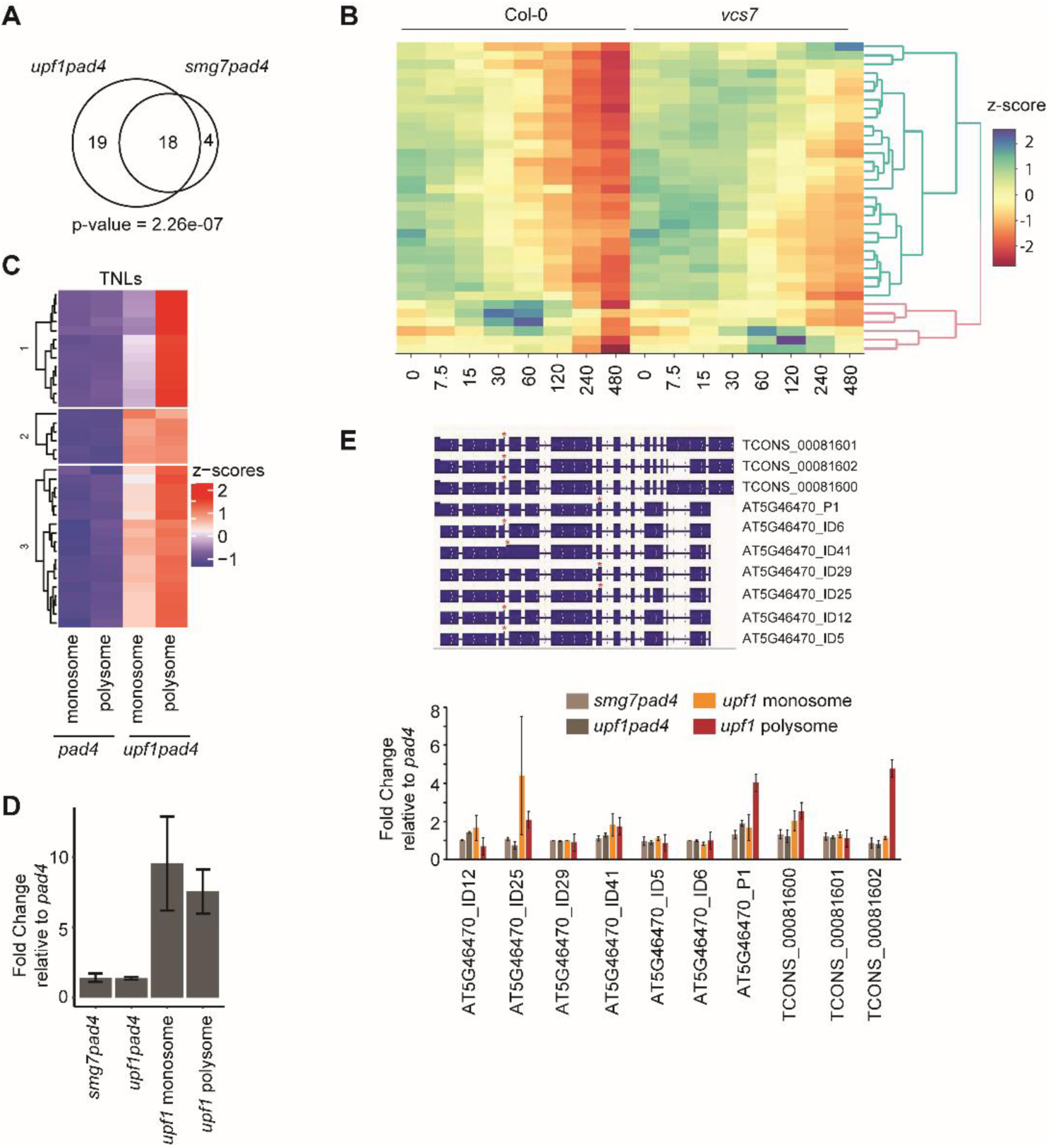
Expression of TNLs in *upf1 pad4* plants. (A) Area proportional Venn diagram depicting overlap of up-regulated {log2 (fold change) ≥ 1, BH-corrected p-value ≤ 0.01 TNL genes in *smg7 pad4* and *upf1 pad4* compared to *pad4*. Statistical significance of the overlap was tested by Fischer’s exact test. (B) Clustered heatmap showing degradation of the NMD-regulated TNL genes over time in wild type and *vcs7* after transcription inhibition. (C) Clustered heatmap showing expression of TNL genes in monosomal and polysomal fractions of *pad4* and *upf1 pad4* plants. (D) Total *RPS6* RNA expression and its abundance in polysomes and monosomes in *smg7 pad4* and *upf1 pad4* plants relative to *pad4*. (E) Expression of RPS6 transcripts and their abundance in polysomes and monosomes in *smg7 pad4* and *upf1 pad4* plants relative to *pad4*. The RPS6 transcript models are shown; stop codon locations are indicated (*). Error bars in (D) and (E) represent standard error of 3 biological replicates.

We next explored our ribosome profiling data to examine the effect of UPF1 on TNL translation. We observed a profound increase in the association of UPF1-regulated TNLs with monosomes and polysomes in *upf1 pad4* compared to the *pad4* control (**Figure 7C**). The enhanced association of TNL RNAs with ribosomes, which is particularly evident for the polysomal fraction, demonstrates that UPF1 acts as a repressor of TNL translation. In our previous study, we showed that RPS6 is one of the TNLs responsible for the increased immunity and growth defects in *smg7* (Gloggnitzer et al. 2014). The *RPS6* locus exhibits typical features of NMD targets, including a long 3’UTR with numerous introns. Curiously, we previously found that the expression of RPS6 in *smg7 pad4* mutants is only 1.6-fold higher than wild type, despite its strong phenotypic effect (Gloggnitzer et al. 2014)(**Figure 7D**). While we confirmed this relatively modest increase in RPS6 RNA in *upf1 pad4* plants, we observed a 9- and 7-fold increase in the association of RPS6 with monosomes and polysomes, respectively (**Figure 7D**). To further explore the effect of UPF1 on the *RPS6* locus, we analyzed individual *RPS6* mRNAs from transcriptome reassembly. We identified three new splice variants in addition to the seven that were already annotated in the AtRTD2 transcriptome (**Figure 7E**). Interestingly, while all transcripts harbored NMD features, only a subset were translationally up-regulated in *upf1 pad4*. Together, these data indicate that, at least for RPS6, UPF1 exerts a much larger effect on translation than on steady-state mRNA levels.

## Discussion

NMD is a molecular pathway consisting of consecutive steps to recognize, validate, and degrade aberrant RNAs. However, recent progress in our understanding of its mechanistic details, particularly in humans, indicates that NMD is far from being a linear pathway. NMD target recognition and degradation can be achieved by independent routes that can be partially redundant or act in different contexts. Moreover, inactivation of different NMD factors has different phenotypic outputs. While knock-out or knock-down of UPF1 and UPF2 is embryonic lethal in flies and zebra fish, inactivation of UPF3 has little effect on viability (Wittkopp et al. 2009; Avery et al. 2011). These observations suggest that NMD is a molecular network integrated around UPF1, rather than a linear pathway. In support of this hypothesis, we provide evidence that different NMD mutants in Arabidopsis show different degrees of NMD efficiency and have different phenotypes. Here we show that UPF3 has a comparatively smaller effect on NMD than UPF1 and SMG7 in Arabidopsis. This is consistent with results obtained in tobacco using virus induced gene silencing assay (Kerenyi et al. 2008) and indicates a less prominent role of the UPF3 pathway in PTC recognition in plants.

Further, we found that inactivation of SMG7 has a substantially smaller effect on the abundance of PTC transcripts and the global transcriptome than UPF1. This observation indicates that *smg7* retains residual NMD activity, which partially explains its less severe phenotypic consequences than *upf1*. The loss of SMG7 may, to some extent, be compensated by increased UPF1 activity as it remains hyperphosphorylated in *smg7* (Kesarwani et al. 2019). Nonetheless, this does not explain how RNA is degraded in *smg7*. In animals, RNA decay is achieved by the partially redundant SMG6 and SMG5/7 pathways (Colombo et al. 2017; Nelson et al. 2018). However, Arabidopsis lacks SMG6 and SMG5 paralogues (Kerenyi et al. 2008; Riehs-Kearnan et al. 2012) and the only Arabidopsis protein related to SMG7 is SMG7L, which does not act in NMD (Benkovics et al. 2011; Capitao et al. 2018). This indicates the existence of alternative mechanism(s) that link UPF1 with RNA degradation. UPF1 was shown to directly interact with the decapping enzyme DCP2 in yeast and human (He and Jacobson 1995; Lykke-Andersen 2002; Loh et al. 2013). A novel UPF1-mediated RNA decay mechanism involving the interaction of the mRNA decapping machinery with phosphorylated UPF1 was described in human cells (Nicholson et al. 2018). An analogous pathway may also operate in plants. RNA tethering experiments showed that UPF1 can trigger SMG7-independent RNA degradation by XRN4, a 5’-3’ exonuclease that acts on decapped transcripts (Merai et al. 2012), and the core components of the mRNA decapping complex co-immunoprecipitated with UPF1 in Arabidopsis (Chicois et al. 2018). Furthermore, transcripts identified in this study that are specifically up-regulated in *upf1 pad4*, but not in *smg7 pad4*, appear to be processed through the decapping pathway (**Figure 5C**). A recent analysis of RNA degradation intermediates that accumulate upon deletion of Arabidopsis XRN4 detected signatures of endonucleolytic cleavage within putative NMD substrates (Nagarajan et al. 2019). This indicates that, in addition to decapping followed by exonucleolytic degradation, plant NMD utilizes yet another RNA decay mechanism that relies on an unknown endonuclease.

Genome-wide studies revealed that NMD affects 5-20% of the transcriptome. However, these studies were performed either with weaker alleles of NMD genes or siRNA-mediated knock downs that retain residual NMD activity (Mendell et al. 2004; Kurihara et al. 2009; Rayson et al. 2012; Drechsel et al. 2013; Hurt et al. 2013; Aliouat et al. 2019; Nagarajan et al. 2019). Here we took advantage of viable *upf1 pad4* plants to assess the full impact of NMD on the Arabidopsis transcriptome. We found deregulation of 21% of expressed genes in *upf1 pad4* using high stringency thresholding, which is substantially more than in *smg7, upf3*, and weaker *upf1* mutants (Kurihara et al. 2009; Rayson et al. 2012; Gloggnitzer et al. 2014). This reinforces the view that NMD plays a central role in transcriptome regulation. Alternative splicing coupled to NMD (AS-NMD) was proposed to be an important mechanism shaping plant transcriptomes (Kalyna et al. 2012; Drechsel et al. 2013). Our data unmask the full extent of this regulation as, at some loci, more than 90% of the splicing events result in PTC+ transcript isoforms, which are efficiently removed by NMD (**Figures 1D and 4B**).

In addition to AS-NMD, our differential transcript analysis indicates that UPF1/NMD directly affects splicing. This is apparent at loci where the abundance of one splice variant switched to another upon UPF1 inactivation (**Figure 4B**). As evident from the GO analysis in *upf1 pad4* mutants, this can be partially due to deregulation of splicing factors and proteins involved in RNA processing, some of which are controlled by AS-NMD (**Figure 1D**)(Kalyna et al. 2012). In addition, UPF1 may be more directly involved in splicing. Studies in human and Drosophila cells showed that UPF1 associates with nascent RNAs co-transcriptionally in the nucleus and this association is required for the release of mRNA from sites of transcription (de Turris et al. 2011; Singh et al. 2019). Thus, UPF1 may directly affect homeostasis of splicing factors on transcribed RNA and influence splicing efficiency. Proteomic data suggesting an association between splicing factors and UPF1 also hint at a more intimate connection between splicing and NMD in Arabidopsis (Sulkowska et al. 2019).

mRNAs processed by NMD become translationally repressed through UPF1-mediated inhibition of translation initiation (Isken et al. 2008). Thus, it is expected that inactivation of NMD will lead to translational derepression of aberrant transcripts. Nevertheless, this question has only been addressed at a genome-wide level in *Saccharomyces cerevisiae* where NMD substrates are less efficiently translated but inactivation of NMD factors did not increase their translation (Celik et al. 2017). Here we used quantification of RNAs in monosomal and polysomal fractions as a proxy of their translational status. Monosomes are traditionally perceived as RNPs with a newly assembled ribosome at the start codon and translationally inactive. However, a recent study in *Saccharomyces cerevisiae* determined that the majority of monosomes are actively translating and contained mainly transcripts with slow rates of translational initiation (Heyer and Moore 2016). We found that a general feature of Arabidopsis transcripts with a high monosome/polysome ratio is their short length (average ∼1000 nt). Because shorter transcripts accommodate fewer ribosomes during translational elongation, this correlation supports the idea that a portion of monosomal transcripts are actively translated also in Arabidopsis.

Intriguingly, inactivation of UPF1 led to a global shift of transcripts to the monosomal fraction. At least two non-exclusive scenarios can explain this observation. First, the absence of NMD was shown to increase the incidence of mRNA-containing transcription errors (Gout et al. 2017). Some of the errors may generate PTCs near the ATG codon, producing short ORFs that would predominantly associate with monosomes. Second, *upf1 pad4* mutants exhibit a suppression of genes involved in translation and ribosome assembly. A shortage of ribosomes and translation factors may reduce the rate of translation initiation and globally decrease the proportion of mRNAs associated with polysomes. The down-regulation of translational processes seems to be specifically associated with inactivation of UPF1, as no effect on these genes was detected in SMG7 or UPF3 deficient plants. The mechanism by which UPF1 downregulates genes involved in translation is unknown, but it may represent a more general phenomenon as suppression of ribosomal proteins was also recently observed in UPF1-depleted human cells (Aliouat et al. 2019).

While the bulk of mRNA shifts to monosomes, we found a subset of transcripts that increase their association with polysomes upon UPF1 deletion, implying an increase in translation. These transcripts are, on average, less stable and abundant, indicating rapid turnover. Increased polysomal association was also observed for putative NMD targets, supporting the idea that NMD acts as a translational repressor of aberrant RNAs. UPF1 has a particularly pronounced impact on translation of TNL immune receptors. TNLs constantly survey for the presence of pathogen proteins within plant cells and their activation results in a strong defense response that may culminate in cell death (Cui et al. 2015). The expression of TNLs is therefore under strict control, as their overexpression triggers autoimmunity that is associated with a massive fitness cost. Hence, there must be regulatory mechanisms that permit the surveillance function of TNLs while limiting their negative impact on fitness (Lai and Eulgem 2018). NMD may represent such a mechanism. We have previously found that a substantial portion of Arabidopsis TNLs are regulated via NMD-mediated RNA turnover and that NMD is down-regulated upon bacterial infection. We showed that this mechanism contributes to plant defense and proposed a role for NMD in the evolution of TNL receptors (Gloggnitzer et al. 2014; Raxwal and Riha 2016). A recent study showed that the infection-induced NMD down-regulation is mediated through UPF1 ubiquitination and degradation (Jung et al. 2020). Here, we demonstrate that inactivation of NMD not only increases the level of TNL transcripts, but it releases their translation, providing a means for a rapid response to pathogen infection. In fact, in the case of RPS6, the effect on translation is much greater than on the steady-state level of mRNA (**Figure 7D**,**E**). The *RPS6* contains a long 3’UTR with multiple introns, making it a strong NMD target. Single molecule experiments in human cells revealed that, on average, only 10% of translation termination events occurring at a PTC trigger NMD (Hoek et al. 2019). This implies that, despite being targeted by NMD, *RPS6* mRNA can still produce sufficient amounts of protein for surveillance, but under the threshold required for triggering immunity. Suppression of NMD upon infection would increase the translation of already present RPS6 transcripts and boost plant defense. Thus, the ability of NMD to efficiently control translation may be more physiologically relevant than its impact on RNA stability.

In conclusion, this study demonstrates a central role for UPF1 in post-transcriptional and translational gene regulation. While the deregulation of gene expression observed in UPF1-null plants can be largely attributed to the loss of NMD, our data indicate effects of UPF1 beyond the degradation of aberrant RNAs and imply a more direct role for UPF1 in regulating splicing and translation.

## Material and methods

### Plant material and growth conditions

The following Arabidopsis mutant lines were used in this study: *upf1-3 pad4-1* and *smg7-1 pad4-1* (Riehs-Kearnan et al. 2012), *upf1-5* (Arciga-Reyes et al. 2006), *upf3-1* (Hori and Watanabe 2005), and *pad4-1* (Jirage et al. 1999). *A. thaliana* accession Columbia-0 was used as a wild-type control. Plants were grown on soil with 16 hr light/8 hr dark photoperiod (21°C, 60% relative humidity) with the exception of *upf1-3 pad4-1* mutants that were germinated on MS media supplemented with 1% sucrose and transferred to soil 10 days after germination.

### High-resolution RT PCR panel

RNA was extracted from leaves of 3-week old plants. RNA extraction and first strand cDNA synthesis were performed as previously described (Gloggnitzer et al. 2014). The high-resolution RT-PCR and splicing analysis were performed according to Simpson et al. (Simpson et al. 2008). The sizes of the RT-PCR products reflect different alternative splicing events providing relative abundance data of transcript isoforms containing the different splicing outcomes. The ratio of peak abundance is compared among different mutants and controls. For each ASE, the distribution of PTC+ transcripts was plotted in R using ggplot2 (Wickham 2009). The Mann-Whitney test was performed in R to distinguish significance differences between NMD mutants.

### Gene expression comparison analysis of NMD mutants

Strand specific libraries were prepared from ribosomal-RNA depleted mRNA using ScriptSeq Complete Plant kit (Epicentre) and sequenced on an Illumina HiSeq2500. For wild type, *upf1-5, upf3-1, pad4*, and *smg7 pad4*, previously published RNA-seq datasets were re-analyzed (GEO datasets GSE41432 and GSE55884) (Drechsel et al. 2013; Gloggnitzer et al. 2014). The expression of transcripts in AtRTD2 (AtRTD2-QUASI) was quantified using Kallisto version 0.43.0 (Bray et al. 2016; Zhang et al. 2017). Subsequently, differential expression analysis was performed using the limma-voom pipeline enabled in the 3D RNA-seq tool (Guo et al. 2019). Low-expressed transcripts were filtered if they did not have ≥1 sample with ≥1 CPM expression. A gene was determined as expressed gene if one of its transcripts was expressed. The resulting expressed transcripts were used to estimate the batch effect among samples and the batch effect was removed from samples using RUVr (Risso et al. 2014). The libraries were length normalized using the trimmed means of M value normalization method and differential expression analysis was performed with limma-voom (Law et al. 2014). 3D RNA-seq identifies significantly DE genes and transcripts, DE and DAS genes and DAS genes in contrast group comparisons (Guo et al. 2019). The heatmap of significantly expressed genes/transcripts was produced in R using Complexheatmap (Gu et al. 2016). The significantly enriched (BH corrected p-value <0.05) gene ontological terms were extracted from the panther database (Ashburner et al. 2000; Mi et al. 2017; Carbon et al. 2018) and plotted using R. The overlap analysis was performed using Vennerable (Swinton 2019).

### Construction of UPF1 reference transcriptome database (UPF1 RTD)

Seedlings (wild type, *pad4* and *upf1 pad4*) were grown in liquid culture (Gamborg’s medium: 3.2 g/l Gamborg’s B5 salts+minimal organics, 1 ml/l 1000×Gamborg’s vitamins, 0.5 g/l MES, 3% sucrose, pH 5.9 and supplemented with 160 µg/ml L-Lysine and 160 µg/ml L-Arginine) in a 250 ml flask (Lewandowska et al. 2013). Briefly, sterile seed were germinated in liquid medium and grown at 22°C in a 16 h light/8 h dark cycle with shaking for 19 days. Medium was exchanged every two days. Total RNA was extracted from the seedlings and libraries were constructed from three independent biological replicates following polyA selection using instructions for TruSeq RNA library preparation (Illumina protocol 15026495 Rev. B). Libraries were sequenced on the Illumina HiSeq2500 platform generating 100 bp paired-end reads. Construction of UPF1 RTD was performed as previously described (Zhang et al. 2017). In brief, sequencing read quality was checked with FastQC followed by trimming of low quality reads (<Q20). Quality-filtered reads were aligned to the Arabidopsis genome using STAR (Dobin et al. 2013) with AtRTD2 as the reference transcriptome. After mapping, transcript models were assembled using two assemblers (Cufflinks and Stringtie) with the GTF guided mode (Trapnell et al. 2012; Pertea et al. 2015). The two transcriptome assemblies (Cufflinks and Stringtie) were merged using cuffmerge (Trapnell et al. 2012) to generate a single assembly. This assembly was passed through additional quality filters as previously described (Zhang et al. 2017). These filters included removal of transcripts with non-canonical splice junctions (not GT..AG, GC..AG, or AT..AC), low expressed transcripts (minimum 1TPM in at least 2 samples) and transcripts with unsupported splice junctions (less than 3 reads in at least 2 samples). Redundant, fragmentary, and chimeric transcripts were removed as previously described (Zhang et al. 2017). The resulting UPF1 RTD contained 83,546 transcripts which were used for further analysis. Isoforms not present in AtRTD2 (Zhang et al. 2017) were reported as novel isoforms. Similarly, transcripts originating from genomic locations other than those in Araport11 (on which the AtRTD2 transcriptome is anchored) were termed as novel loci.

### PTC definition

To overcome issues with mis-annotation of ORFs, transcripts were translated with TranSuite (JCE, RZ, JB unpublished). Briefly, each isoform of a gene was translated to identify the longest reading frame. The start codon of the isoform with the longest reading frame was fixed as the +1 codon for the rest of the isoforms for further translation. Stop codons were defined as premature termination codons if they were present ≥50 nt upstream of an exon-exon junction.

### Polysome profiling

The monosome and polysome fractions were isolated from leaves of soil grown plants using a published protocol (Juntawong et al. 2015) with minor modifications. In brief, 150 mg of leaf tissue was ground in liquid nitrogen and transferred to 500 µl of polysome extraction buffer (200 mM Tris-Cl pH 9.0, 200 mM KCL, 25 mM EGTA, 35 mM MgCl2, 1X detergent mix, 1% PTE, 5 mM DTT, 1 mM PMSF, 50 µg/ml cycloheximide, 50 µg/ml chloramphenicol, 1X protease inhibitor, RNase Out). The solution was centrifuged at 14,000 rpm for 15 min and the supernatant was loaded on a 15-60% sucrose gradient. The gradient was ultracentrifuged at 50,000 rpm for 65 min, fractionated and RNA content in the fractions measured and visualized to identify monosomal and polysomal fractions. The RNA was extracted from monosomal and polysomal fractions as previously described (Mustroph et al. 2009). Strand specific libraries were prepared from ribosomal-RNA depleted mRNA using ScriptSeq Complete Plant kit (Epicentre) and sequenced on the Illumina HiSeq2500 platform (100 bp single-end reads). Transcripts were quantified using the AtRTD2 and Kallisto (version 0.43.0; (Bray et al. 2016)) followed by differential expression analysis using limma-voom pipeline in 3D-RNAseq (Guo et al. 2019). The polysome abundance score for each gene was calculated as [gene expression (TPM) in polysome / gene expression in monosome + polysome (TPM)]. The polysome abundance score ranges from 0 to 1 with, higher values representing more mRNA in polysomes than monosmes. The density kernel estimation of genes in *pad4* and *upf1 pad4* was calculated based on the polysome abundance score. Tests for significant differences in the two-density plot was calculated using the Kolmogorov-Smirnov test.

The difference between the polysome abundance score for each gene in *pad4* and *upf1 pad4* sample was calculated as the polysome abundance score of *upf1 pad4* vs. *pad4*. The monosome:polysome score was calculated as fold enrichment between monosomal and polysomal bound RNA [gene expression in monosomes (TPM)/ gene expression in polysomes (TPM)] and the test for significant differences was calculated using the limma-voom pipeline.

### RNA decay analysis

Raw sequencing data from RNA decay time series (0 to 480 minute) of Col-0 and *vcs7* was downloaded from the Gene Expression Omnibus (GEO) database (accession no. GSE86361). The reads were used to quantify gene expression of the AtRTD2 assembly using Kallisto and the lima-voom pipeline enabled in 3D-RNAseq. The relative fold change for each gene, on a log 2 scale, was calculated for each time point with respect to control (0 minute). The heatmap was plotted using heatmaply (Galili et al. 2018) after scaling the rows to Z-score.

### Real time PCR

Total RNA was extracted from 3-week old soil grown plants (*pad4, smg7pad4*, and *upf1pad4*) as previously described (Gloggnitzer et al. 2014). DNA was removed from total RNA using Turbo DNase in accordance to the manufacturer’s protocol. cDNA was prepared from DNase treated total RNA (2.5 µg) using SuperScript IV Reverse Transcriptase. qPCR was performed using Kappa SYBR Fast qPCR master mix on a Light Cycler 96. cDNA was calibrated with AT4G26410 and the ΔΔCt method (Livak and Schmittgen 2001). The list of primers used are presented in **Supplemental Table 6**.

## Acknowledgements

We thank to Zuzana Feketová for assistance with polysome preparation. This research was supported by the Ministry of Education, Youth and Sports of the Czech Republic, European Regional Development Fund-Project “REMAP” (No. CZ.02.1.01/0.0/0.0/15_003/0000479 to KR), the Czech Science Foundation (16-18578S to KR), Biotechnology and Biological Sciences Research Council (BBSRC; BB/P009751/1 to JB); Scottish Government Rural and Environment Science and Analytical Services division (RESAS to CS, JB and RZ), and the BBSRC EASTBIO PhD studentships (to JCE).

