## Supplemental Figures for "Nonsense mediated RNA decay factor UPF1 is critical for post-transcriptional and translational gene regulation in Arabidopsis"

### Supplemental Material

Supplemental Figure 1

Supplemental Figure 2

Supplemental Figure 1

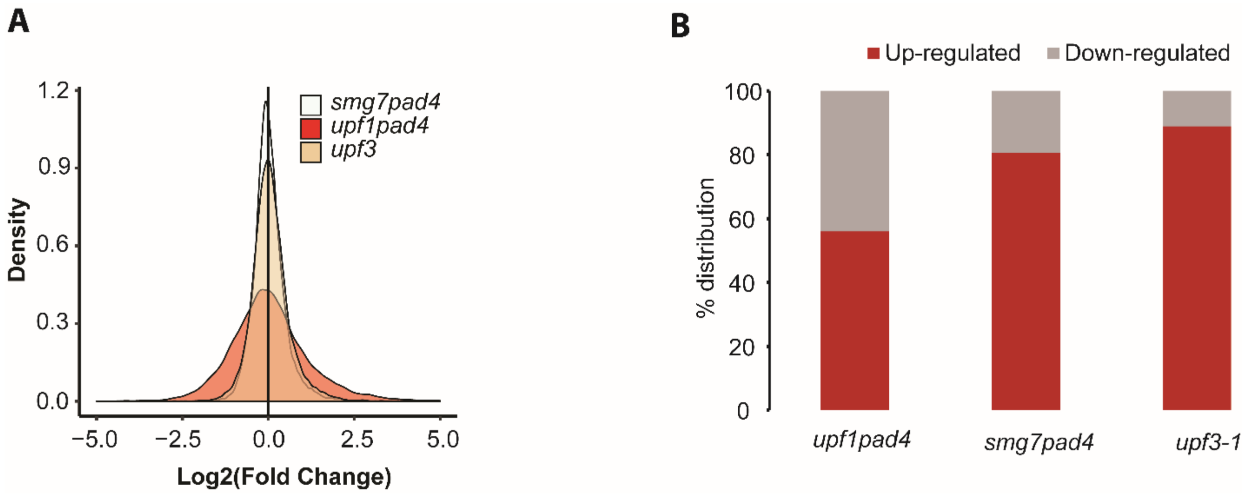


**Supplemental Figure 1. Differential gene expression in transcriptomes of *upf1 pad4*, *smg7 pad4*. and *upf3* mutants.** (A) Smoothed Kernel density estimate of gene expression showing a Gaussian distribution in indicated mutants compared to their respective controls (*pad4* or wild type). (B) Bar plot showing the percentage distribution of up-regulated and down-regulated genes. Absolute number of DE genes: *upf1 pad4* – 3623, *smg7 pad4* – 612, *upf3* – 279.

Supplemental figure 2

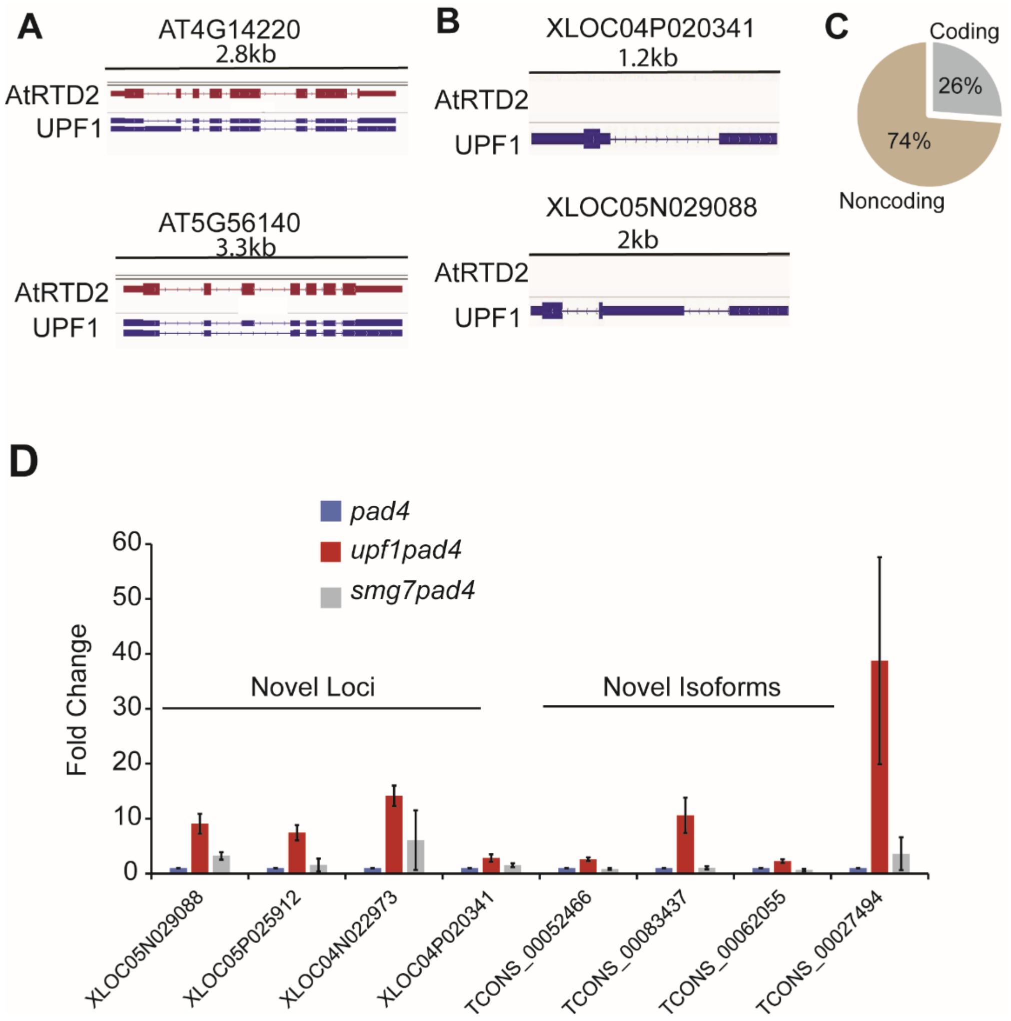


**Supplemental Figure 2. Novel transcripts detected in *upf1 pad4* transcriptome.** IGV genome browser image depicting two examples of transcript models for novel isoforms (A) and novel loci (B) in reference (TAIR10, red color) and UPF1 assembly (blue color). (C) Pie chart representing the percentage distribution of coding (grey) and non-coding (red) transcripts in novel loci. (D) qRT-PCR analysis of selected novel isoforms and novel loci in *smg7 pad4* and *upf1 pad4* relative to *pad4*. Error bars represent standard error of three biological replicates.
